# Can Orthopaedics become the Gold Standard for Reproducibility? A Roadmap to Success

**DOI:** 10.1101/715144

**Authors:** Ian A. Fladie, Sheridan Evans, Jake Checketts, Daniel Tritz, Brent Norris, Matt Vassar

## Abstract

**Background:** Scientific research is replete with poor accessibility to data, materials, and protocol, which limits the reproducibility of a study. Transparency with regard to materials, protocols, and raw data sets enhances reproducibility by providing the critical information necessary to verify, replicate, and resynthesize research findings. The extent to which transparency and reproducibility exist in the field of orthopaedics is unclear. In our study, we aimed to evaluate transparency and reproducibility-related characteristics of randomly sampled publications in orthopaedic journals.

**Methods:** We used the National Library of Medicine catalog to identify English language and MEDLINE-indexed orthopaedic journals. From the 74 journals meeting our inclusion criteria, we randomly sampled 300 publications using a refined PubMed search that were published between January 1, 2014, and December 31, 2018. Two investigators were trained for data extraction and analysis. Both investigators were blinded and independently extracted data from the 300 studies.

**Results:** Our initial search yielded 68,102 publications, from which we drew a random sample of 300 publications. Of these 300 publications, 286 were screened for empirical data and 14 were inaccessible. For analysis purposes, we excluded publications without empirical data. Of the 182 with empirical data, 13 studies (7.1%) included a data availability statement, 9 (4.9%) reported materials were available, none (0.0%) provided analysis scripts, 2 (1.1%) provided access to the protocol used, 5 (2.7%) were preregistered, and only 2 (1.1%) provided a statement about being a replicated study.

**Conclusions:** Components necessary for reproducibility are lacking in orthopaedic surgery journals. The vast majority of publications did not provide data or material availability statements, protocols, or analysis scripts, and had no preregistration statements. Intervention is needed to improve reproducibility in the field of orthopaedics. The current state of reproducibility in orthopaedic surgery could be improved by combined efforts from funding agencies, authors, peer reviewers, and journals alike.

**Level of Evidence:** N/A

## Introduction

The National Institutes of Health (NIH) considers research rigor and reproducibility to be integral parts of modern research practice^1^. Reproducibility—the ability to conduct a study using the same materials and protocol to obtain similar results—is the foundation of verifying and improving scientific practice^2^. Transparency with regard to materials, protocols, and raw data sets used to conduct original research enhances reproducibility by providing the means to verify, replicate, and resynthesize findings of well-established literature^3^. If a study is reproducible, the trial design could be disseminated among various specialties and benefit patients in all fields of medicine, including orthopaedics. Despite the benefits of reproducible studies, many cases of studies providing limited access to data have been documented. For example, in an analysis of 441 studies, Iqbal et al. discovered that approximately 66% of the studies provided empirical data and none provided access to any raw data^4^. Furthermore, a survey of 1,576 researchers found that approximately 90% of respondents believe there is a crisis in research reproducibility^5^. The field of orthopaedics is no exception, as illustrated by a report by Beard et al., who found in a comparison of subacromial decompression, arthroscopy, and no surgical treatment that the results were insignificant^6^.

Scientific research is replete with poor accessibility to data, materials, and protocols, which limits reproducibility. In an analysis of 50 high-profile papers, the Reproducibility Project in Cancer Biology found that only 18 had sufficient materials to replicate the findings^7^. Additionally, in 2015 an open collaboration to replicate 100 studies from 3 peer-reviewed psychology journals found that only 36% of the results remained statistically significant, despite 97% of the original findings being statistically significant^8^. Such findings have prompted funding agencies, journals, and other research stakeholders to enact policies and procedures toward improving reproducibility. For example, the NIH established the Rigor and Reproducibility Initiative, which provides guidance for authors to enhance reproducibility through a focus on the scientific premise, scientific rigor, biological variables, and authentication of results^9^. Journals have also enacted policy changes to promote reproducible and transparent research practices. For example, in a study of 21 orthopaedic surgery journals, Checketts et al. reported that 52% required clinical trial registration^10^. Furthermore, several studies have examined the implementation of data-sharing policies, finding that their enactment led to an increase in the publications that included raw data^11,12^. With publication retractions in orthopaedics increasing, primarily due to academic misconduct and fraud^13^, the need to verify trials by reproducing studies has become more urgent. In this study, our primary objective was to evaluate publications in the field of orthopaedic surgery, using the 8 indicators of research transparency and reproducibility defined by Hardwicke et al^13^. Our results highlight areas with the greatest need for improvement and establish a baseline for future investigations.

## Materials and Methods

This cross-sectional study used the methodology of Hardwicke et al., with modifications mentioned later.^13^ The following information is provided on Open Science Framework (https://osf.io/x24n3/): the original training session recording, protocol, raw data, and other pertinent materials.

### Journal and Publication Selection

On May 29, 2019, one of us (D.T.) searched the National Library of Medicine (NLM) catalog for all journals using the subject terms tag “Orthopedics[ST]”. The inclusion criteria (English language and MEDLINE indexed) were then applied to the list of journals. The included journals had their electronic ISSN number (or linking ISSN if electronic was unavailable) extracted. Any additional journals that did not provide full-text publications in English were excluded. The final list of journals was used to search PubMed on May 31, 2019, for all publications. We included publications from January 1, 2014, to December 31, 2018.

### Extraction Training

Prior to data extraction, two investigators (I.F., S.E.) completed training sessions centered on the protocol, study design, and the Google form used for this study. They then separately extracted data from 2 sample publications and met to reconcile differences. Next, they extracted data from 10 additional publications and had another consensus meeting to ensure reliability and accuracy of data extraction. Following extraction, the 2 investigators met for a final consensus meeting to resolve any discrepancies, concluding the training sessions for data extraction.

### Data Extraction

After completing training, two investigators (I.F. and S.E.) extracted data from the 300 sampled publications between June 3, 2019, and June 10, 2019, using a previously tested Google form used during extraction training. After completing data extraction in a duplicate, blinded fashion, investigators met to reconcile disagreements. A third investigator (D.T.) who was available for adjudication was not needed.

### Assessment of Reproducibility and Transparency Characteristics

We used the methodology of Hardwicke et al.^13^ for analyses of transparency and reproducibility of research, with modifications. Full publications were examined for funding disclosures, conflicts of interest, and available materials, data, protocols, and analysis scripts. Publications were coded to fit 2 criteria: no empirical data and studies with empirical data. Publications without empirical data (e.g., editorials, reviews, news, simulations, or commentaries without reanalysis) were only analyzed for statements including conflict of interest, open access, and funding because protocols, data sets, and reproducibility were not relevant. Case studies and case series were listed as empirical studies; however, questions pertaining to the availability of materials, data, protocol, and registration were excluded due to previous study recommendations^14^. Data extraction criteria for each included and excluded study design are described in Table 1.

**Table 1:**
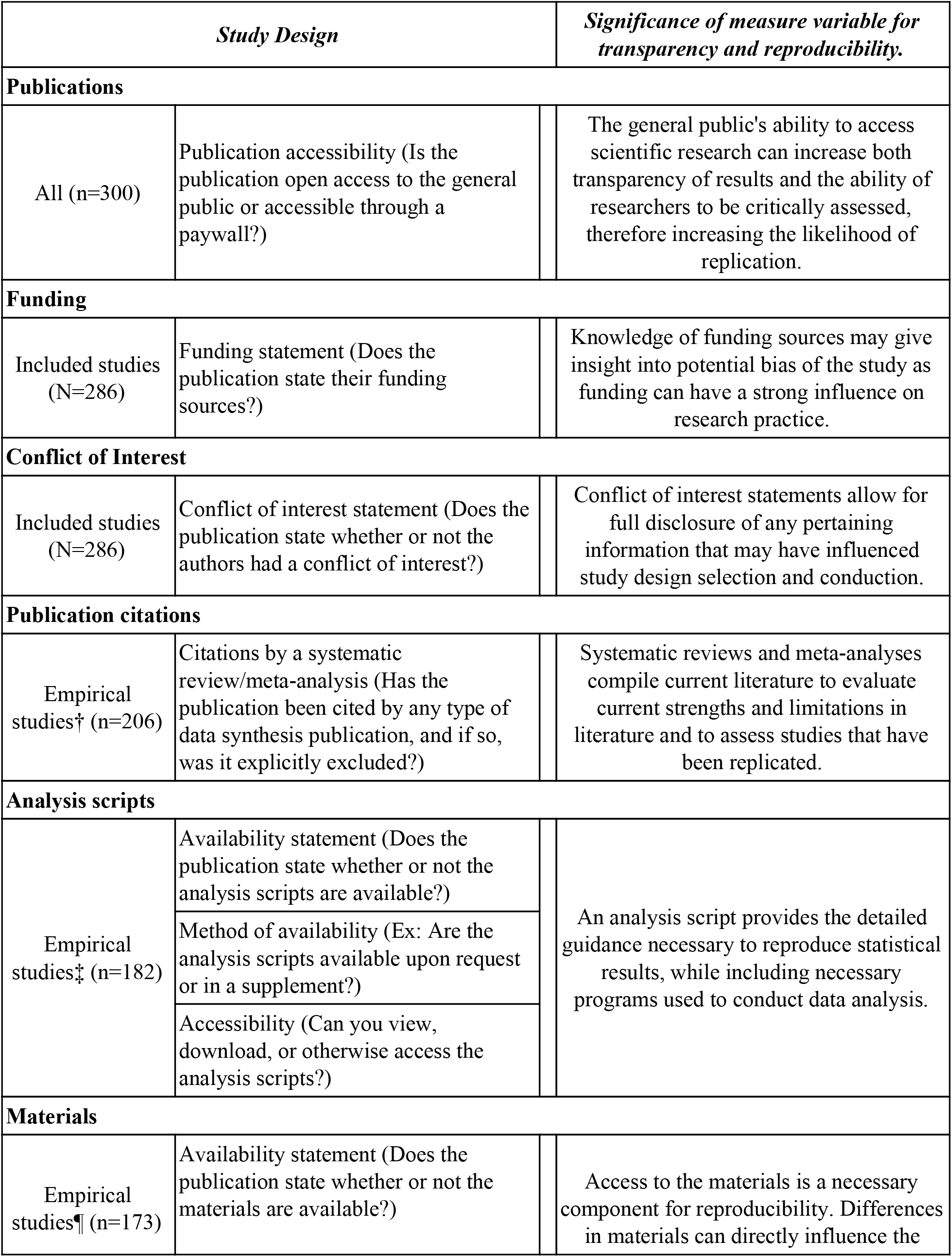
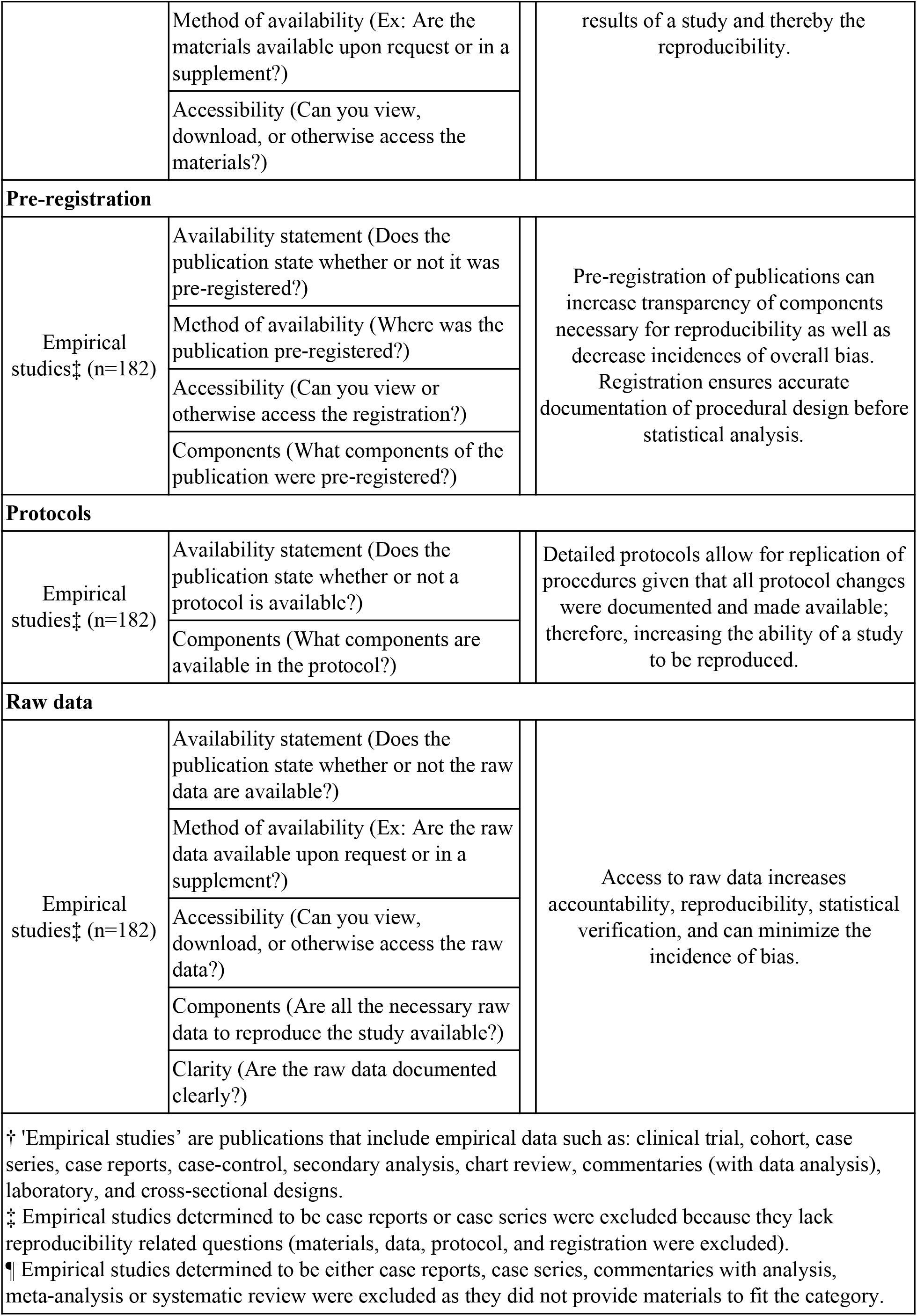
Reproducibility related characteristics. Full detailed protocol outlining each variable measured is available online (https://osf.io/x24n3/)

| Study Design |  | Significance of measure variable for transparency and reproducibility. |
| --- | --- | --- |
| Publications |  |  |
| All (n=300) | Publication accessibility (Is the publication open access to the general public or accessible through a paywall?) | The general public's ability to access scientific research can increase both transparency of results and the ability of researchers to be critically assessed, therefore increasing the likelihood of replication. |
| Funding |  |  |
| Included studies (N=286) | Funding statement (Does the publication state their funding sources?) | Knowledge of funding sources may give insight into potential bias of the study as funding can have a strong influence on research practice. |
| Conflict of Interest |  |  |
| Included studies (N=286) | Conflict of interest statement (Does the publication state whether or not the authors had a conflict of interest?) | Conflict of interest statements allow for full disclosure of any pertaining information that may have influenced study design selection and conduction. |
| Publication citations |  |  |
| Empirical studies† (n=206) | Citations by a systematic review/meta-analysis (Has the publication been cited by any type of data synthesis publication, and if so, was it explicitly excluded?) | Systematic reviews and meta-analyses compile current literature to evaluate current strengths and limitations in literature and to assess studies that have been replicated. |
| Analysis scripts |  |  |
| Empirical studies‡ (n=182) | Availability statement (Does the publication state whether or not the analysis scripts are available?) | An analysis script provides the detailed guidance necessary to reproduce statistical results, while including necessary programs used to conduct data analysis. |
|  | Method of availability (Ex: Are the analysis scripts available upon request or in a supplement?) |  |
|  | Accessibility (Can you view, download, or otherwise access the analysis scripts?) |  |
| Materials |  |  |
| Empirical studies¶ (n=173) | Availability statement (Does the publication state whether or not the materials are available?) | Access to the materials is a necessary component for reproducibility. Differences in materials can directly influence the |
|  | Method of availability (Ex: Are the materials available upon request or in a supplement?) | results of a study and thereby the reproducibility. |
|  | Accessibility (Can you view, download, or otherwise access the materials?) |  |
| Pre-registration |  |  |
| Empirical studies‡ (n=182) | Availability statement (Does the publication state whether or not it was pre-registered?) | Pre-registration of publications can increase transparency of components necessary for reproducibility as well as decrease incidences of overall bias.<br>Registration ensures accurate documentation of procedural design before statistical analysis. |
|  | Method of availability (Where was the publication pre-registered?) |  |
|  | Accessibility (Can you view or otherwise access the registration?) |  |
|  | Components (What components of the publication were pre-registered?) |  |
| Protocols |  |  |
| Empirical studies‡ (n=182) | Availability statement (Does the publication state whether or not a protocol is available?) | Detailed protocols allow for replication of procedures given that all protocol changes were documented and made available; therefore, increasing the ability of a study to be reproduced. |
|  | Components (What components are available in the protocol?) |  |
| Raw data |  |  |
| Empirical studies‡ (n=182) | Availability statement (Does the publication state whether or not the raw data are available?) | Access to raw data increases accountability, reproducibility, statistical verification, and can minimize the incidence of bias. |
|  | Method of availability (Ex: Are the raw data available upon request or in a supplement?) |  |
|  | Accessibility (Can you view, download, or otherwise access the raw data?) |  |
|  | Components (Are all the necessary raw data to reproduce the study available?) |  |
|  | Clarity (Are the raw data documented clearly?) |  |
| † 'Empirical studies' are publications that include empirical data such as: clinical trial, cohort, case series, case reports, case-control, secondary analysis, chart review, commentaries (with data analysis), laboratory, and cross-sectional designs.<br>‡ Empirical studies determined to be case reports or case series were excluded because they lack reproducibility related questions (materials, data, protocol, and registration were excluded). |  |  |

¶ Empirical studies determined to be either case reports, case series, commentaries with analysis, meta-analysis or systematic review were excluded as they did not provide materials to fit the category.

### Publication Citations Included in Research Synthesis and Replication

For both empirical and nonempirical studies, we measured the impact factor of each journal by searching the publication title on Web of Science (https://webofknowledge.com). For empirical studies, we used Web of Science to determine whether studies in our sample were cited in either a meta-analysis, systematic review, or a replication study.

### Assessment of Open Access

Important core components of publications necessary for reproducibility are only available within the full text of a publication. To determine public access to the full text of each publication in our sample, we used Open Access Button (https://openaccessbutton.org), Google, and PubMed. First, we searched the title and DOI using Open Access Button to determine if the publication was available for public access. If the button returned no results or had an error, we searched the publication title on Google, PubMed, and reviewed the journal website to determine if the publication was available without a paywall.

### Modifications

Our study contained the following modifications from Hardwicke et al.^13^. We added the 5-year impact factor and the most recent yearly impact factor we could find (rather than that of a specific year) to our Google form. We also expanded study design options (e.g., cohort, case series, secondary analysis, chart review, and cross-sectional) and included more specific funding options (e.g., university, hospital, public, private/industry, nonprofit, or mixed).

### Statistical Analysis

We used Microsoft Excel to record statistics for each category of our analysis. In particular, we used Excel functions to calculate our study characteristics and results with 95% confidence intervals (95% CIs).

## Results

### Journal and Publication Selection

After searching the NLM catalog, we identified 130 orthopaedic journals. Based on inclusion criteria, 81 journals were selected for our study. The 81 journals yielded 68,102 publications within our time frame, and we randomly selected 300 publications for our sample. Fourteen publications were inaccessible, leaving 286 for analysis. Of the 286 publications, 71 contained no empirical data and were consequently excluded because they lacked reproducibility characteristics. From the remaining 215 publications with empirical data, we excluded 33 that were case studies and case series because these study types are irreproducible. Our final analysis included 182 publications with reproducibility-related characteristics (Figure 1).

**Figure 1:**
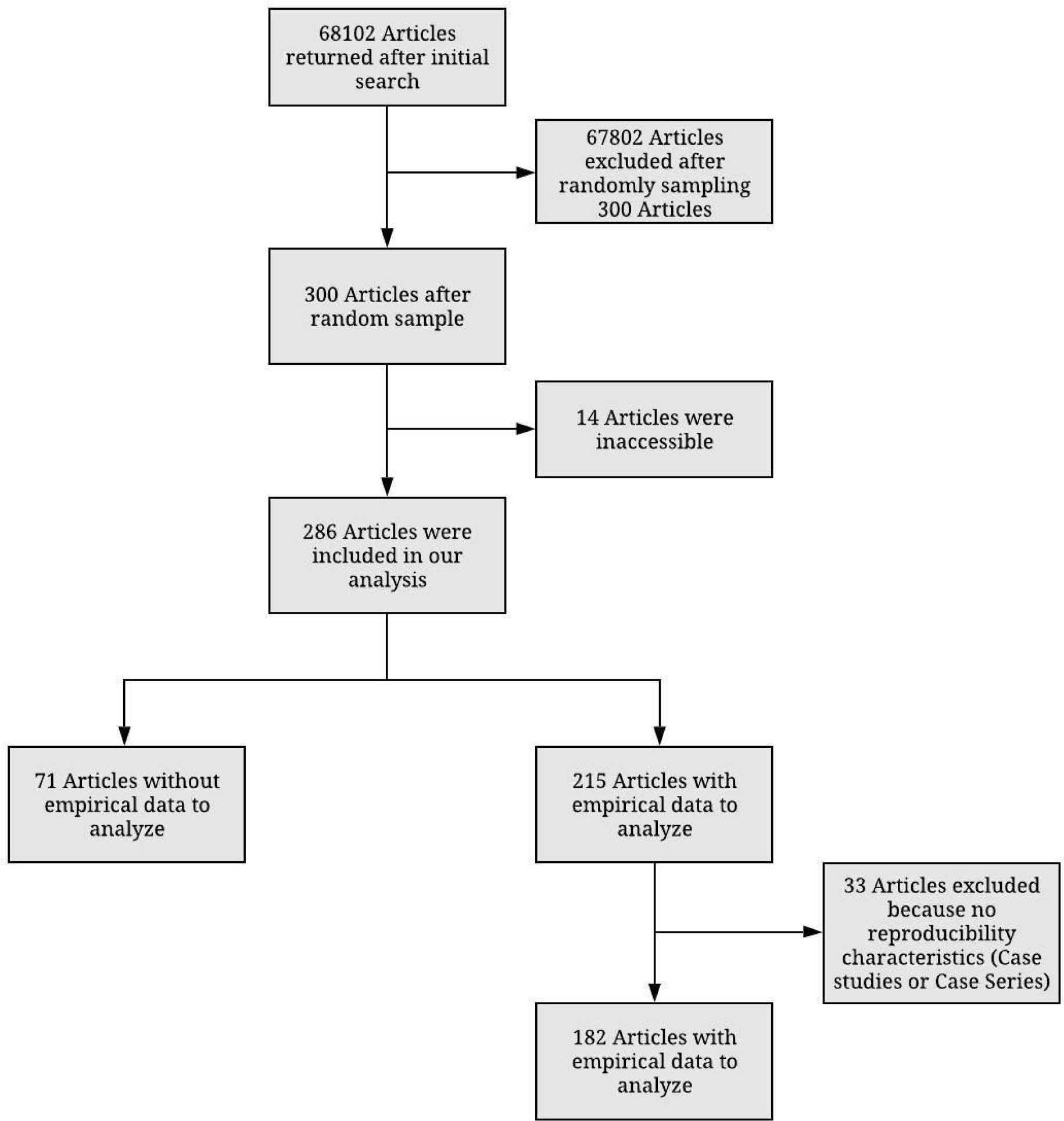
Flow diagram of included and excluded studies in orthopaedic surgery journals

### Sample Characteristics

Of the 286 publications, 32 did not have an available impact factor and the median 5-year impact factor of the other 254 was 2.858 (interquartile range [IQR] 2.108-3.533; Table 3). The majority of corresponding authors and journals were based in the United States. Among the 300 randomly sampled publications, 103 (34.3%) were available to the public according to Open Access Button, 22 (7.3%) were open access and found via other means, and 175 (58.3%) were paywalled. Of the 286 publications that we obtained, 38 (13.3%) contained no statement regarding conflicts of interest, while the majority (248, 86.7%) included a conflict of interest statement. With regard to funding characteristics, approximately half of the 286 publications (140, 49.0%) did not provide any statement about funding. The majority of subject populations were humans, representing 188 (65.7%) of the publications. Additional sample characteristics are presented in Table 2.

**Table 2:** Reproducibility indicators of analyzed orthopaedic publications

| *Characteristics* |  | *Variables* |  |
| --- | --- | --- | --- |
|  |  | **No. (%)** | **95% CI** |
| **Funding (n=286)** | University | 4 (1.40) | - |
|  | Hospital | 1 (0.35) | - |
|  | Public | 34 (11.89) | - |
|  | Private/Industry | 11 (3.85) | - |
|  | Non-profit | 4 (1.40) | - |
|  | No statement listed | 140 (48.95) | - |
|  | No funding received | 71 (24.83) | - |
|  | Mixed funding received | 21 (7.34) | - |
| **Conflict of Interest statement (n=286)** | Statement, one or more conflicts of interest | 59 (20.63) | 16.05-25.21 |
|  | Statement, no conflict of interest | 189 (66.08) | 60.73-71.44 |
|  | No conflict of interest statement | 38 (13.29) | 9.45-17.13 |
| **Data availability (n=182)** | Statement, some data are available | 13 (7.14) | 4.23-10.06 |
|  | Statement, data are not available | 3 (1.65) | 0.21-3.09 |
|  | No data availability statement | 166 (91.21) | 88.00-94.41 |
| **Material availability (n=173)** | Statement, some materials are available | 9 (5.20) | 2.69-7.72 |
|  | Statement, materials are not available | 3 (1.73) | 0.26-3.21 |
|  | No materials availability statement | 161 (93.06) | 90.19-95.94 |
| **Protocol availability (n=182)** | Protocol available | 2 (1.10) | 0.08-2.28 |
|  | No protocol | 180 (98.90) | 97.72-100.08 |
| **Analysis script availability (n=182)** | Statement, some analysis scripts are available | 0 | 0 |
|  | Statement, analysis scripts are not available | 0 | 0 |
|  | No analysis script availability statement | 182 | 1 |
| **Replication studies (n=182)** | Replication | 2 (1.10) | 0.08-2.28 |
|  | No clear statement regarding replication | 180 (98.90) | 97.72-100.08 |
| **Cited by systematic review/Meta-analysis (n=206) †** | No citations | 177 (85.92) | 81.99-89.86 |
|  | Single citation | 22 (10.68) | 7.18-14.17 |
|  | One to five citations | 6 (2.91) | 0.10-4.82 |
|  | Greater than five citations | 1 (0.49) | 0.03-1.27 |
| **Cited by a replication study (n=206)** | No citations | 205 (99.51) | 98.73-100.30 |
|  | Single citation | 1 (0.49) | 0.30-1.27 |
| **Pre-registration (n=182)** | Statement, says was pre-registration | 6 (3.30) | 1.28-5.32 |
|  | Statement, was not pre-registration | 2 (1.10) | 0.01-2.28 |
|  | No, there is no pre-registration statement | 174 (95.60) | 93.28-97.92 |
| **Open access (n=300)** | Yes, found via Open Access Button | 103 (34.33) | 28.96-39.71 |
|  | Yes, found article via other means | 22 (7.33) | 4.38-10.28 |
|  | Could not access through paywall | 175 (58.33) | 52.75-63.91 |
| Abbreviations: CI, Confidence Interval. |  |  |  |
| † No studies were explicitly excluded from the systematic reviews or meta-analysis that cited the original article. |  |  |  |

**Table 3:**
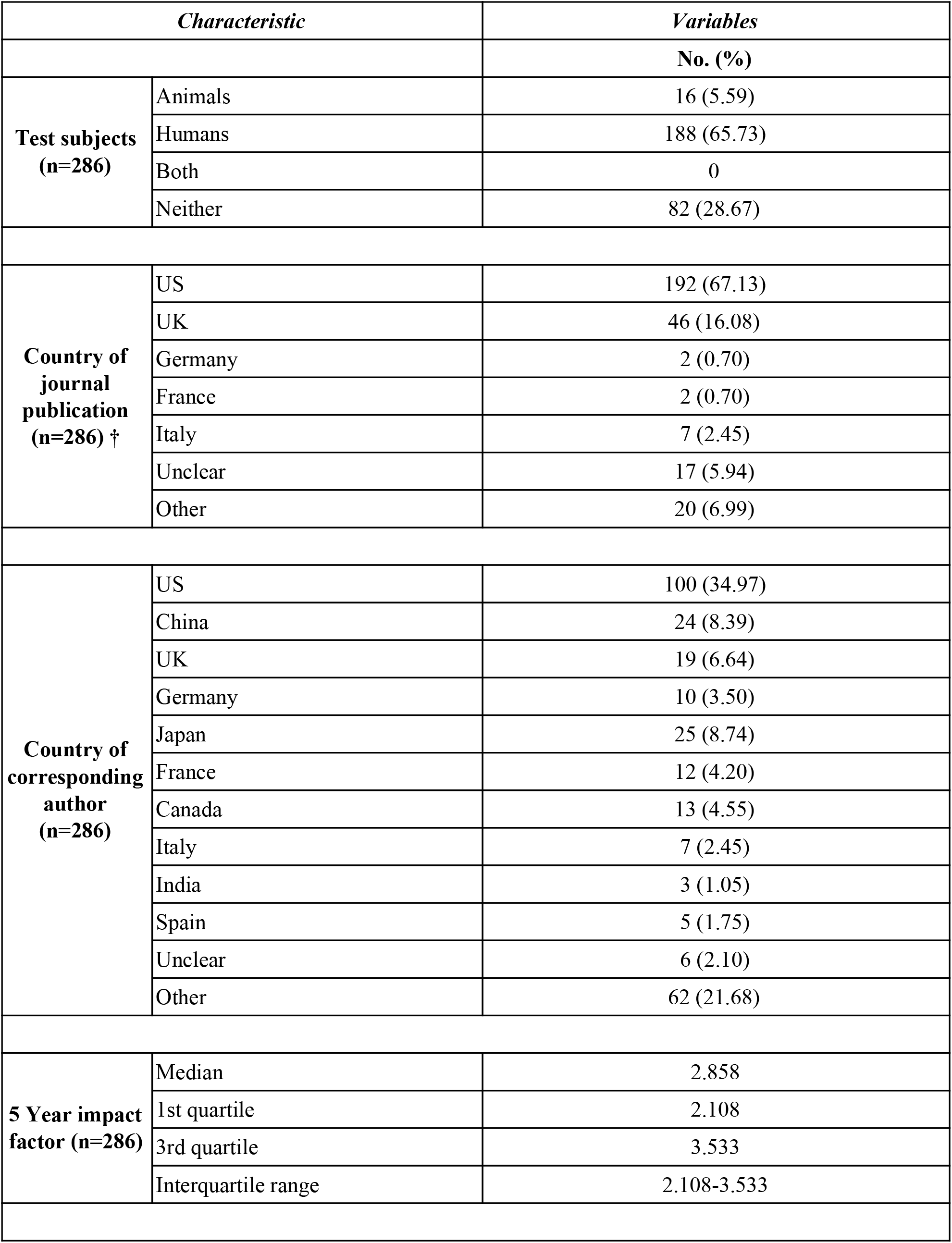
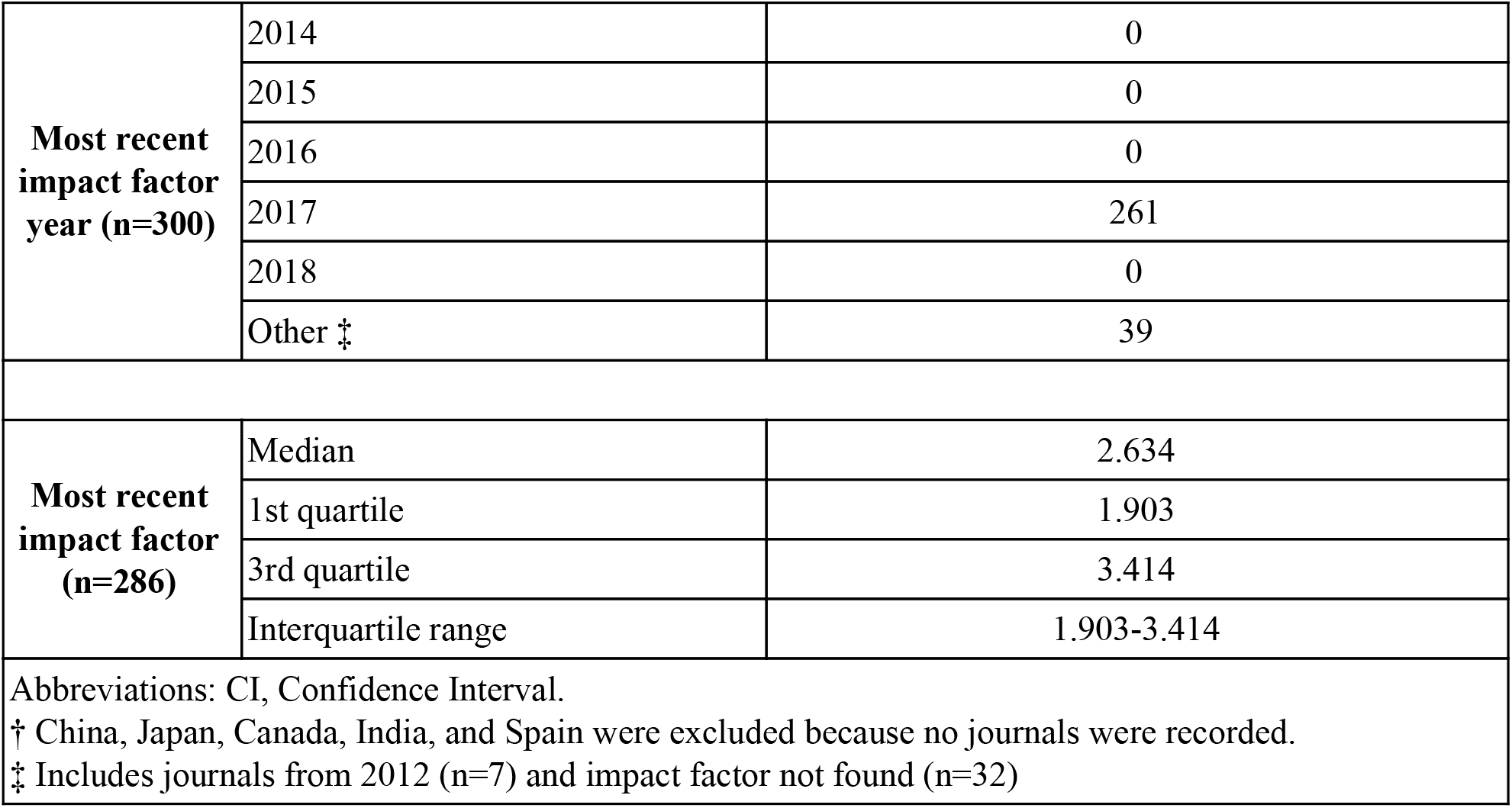
Sample characteristics of analyzed orthopaedic publications

| *Characteristic* |  | *Variables* |
| --- | --- | --- |
|  |  | **No. (%)** |
| **Test subjects**<br>**(n=286)** | Animals | 16 (5.59) |
|  | Humans | 188 (65.73) |
|  | Both | 0 |
|  | Neither | 82 (28.67) |
| **Country of**<br>**journal**<br>**publication**<br>**(n=286) †** | US | 192 (67.13) |
|  | UK | 46 (16.08) |
|  | Germany | 2 (0.70) |
|  | France | 2 (0.70) |
|  | Italy | 7 (2.45) |
|  | Unclear | 17 (5.94) |
|  | Other | 20 (6.99) |
| **Country of**<br>**corresponding**<br>**author**<br>**(n=286)** | US | 100 (34.97) |
|  | China | 24 (8.39) |
|  | UK | 19 (6.64) |
|  | Germany | 10 (3.50) |
|  | Japan | 25 (8.74) |
|  | France | 12 (4.20) |
|  | Canada | 13 (4.55) |
|  | Italy | 7 (2.45) |
|  | India | 3 (1.05) |
|  | Spain | 5 (1.75) |
|  | Unclear | 6 (2.10) |
|  | Other | 62 (21.68) |
| **5 Year impact**<br>**factor (n=286)** | Median | 2.858 |
|  | 1st quartile | 2.108 |
|  | 3rd quartile | 3.533 |
|  | Interquartile range | 2.108-3.533 |
| **Most recent impact factor year (n=300)** | 2014 | 0 |
|  | 2015 | 0 |
|  | 2016 | 0 |
|  | 2017 | 261 |
|  | 2018 | 0 |
|  | Other ‡ | 39 |
| **Most recent impact factor (n=286)** | Median | 2.634 |
|  | 1st quartile | 1.903 |
|  | 3rd quartile | 3.414 |
|  | Interquartile range | 1.903-3.414 |
| Abbreviations: CI, Confidence Interval.<br>† China, Japan, Canada, India, and Spain were excluded because no journals were recorded.<br>‡ Includes journals from 2012 (n=7) and impact factor not found (n=32) |  |  |

### Reproducibility-Related Characteristics

Among the included 182 publications with empirical data, 174 (95.6%) were not preregistered. Eight publications provided preregistration statements; however, only 5 (2.7%) were preregistered, and 3 (1.6%) contained statements that they were not preregistered. Among the 182 publications with empirical data, only 16 (8.8%) provided a data availability statement, with only 13 available data sets provided. Of the data sets available, 5 of the 13 (38.5%) could be accessed and downloaded and only 2 of these 5 (40.0%) provided all raw data to reproduce findings (Supplemental Table 1). Of the 182 publications with empirical data, none provided an analysis script and only 2 (1.1%) provided access to the protocol used to conduct the study. Of the 173 publications that contained empirical data (excluding meta-analyses and commentaries with analysis), 161 (93.6%) did not contain a material availability statement. Of the remaining 12 publications that provided a material availability statements, 3 (25.0%) provided a statement indicating that materials were not available, and 9 (75.0%) provided journal links to material. Additional reproducibility characteristics are shown in Table 2 and Supplemental Table 1.

## Discussion

Our analysis comprised nearly 300 publications in orthopaedic surgery journals. Of the 182 publications analyzed, over 90% of publications did not include a statement of material or data availability, and 95.6% were not prospectively registered. Because the vast majority of the publications we analyzed did not include the necessary resources to reproduce a study, doing so would be challenging at best. Recent research within orthopaedics has consistently shown that improvement is needed to increase transparency and reproducibility and to reduce bias within the field^10,15,16^. Similar research on reproducibility and transparency in other fields has demonstrated methodological shortcomings. Because reproducibility is generally called for to improve biomedical research and no field is superior in this regard, we offer a roadmap for orthopaedics to become the gold standard in producing transparent and reproducible research.

Reproducibility has become a prominent topic in efforts to improve scientific literature and optimize the time and resources spent on research. As such, many recent publications have focused on this topic. Hardwicke et al.^13^ evaluated the reproducibility in 198 social sciences studies and reported findings similar to ours. They found that 84% of publications did not state the availability of materials, 92% lacked a statement regarding data availability, and none of the publications were prospectively registered. Iqbal et al. evaluated the same outcomes in biomedical literature and found that of 268 papers, 99.6% of the publications lacked statements regarding access to materials, none provided a statement regarding access to data, and just 1.9% were prospectively registered^4^. Other authors have evaluated the reproducibility of biomedical research more directly by attempting to reproduce the findings of published studies. Focusing on studies in *Nature* and *Science,* Camerer et al. failed to replicate the results of more than a third and found significantly weaker evidence for the remainder compared with the original reports^17^. Similarly, Aarts et al. found that although 97% of original studies in their sample had statistically significant results, just 36% of the replicated studies met this threshold. Additionally, just 47% of original effect sizes were within the 95% confidence interval in the replication, and only 39% of the measured effects were considered to have replicated the original results. Because measures that increase the ease of reproducing scientific research, such as data and material availability and prospective registration, were not undertaken by the studies in our sample, efforts to reproduce results from the orthopaedic literature could reasonably be expected to demonstrate outcomes similar to those for other fields.

Overall, a crisis in reproducibility clearly affects scientific research as a whole. Orthopaedic research cannot fully help surgeons or patients unless the associated materials and methods are made accessible^18^. Authors have proposed numerous strategies to address this crisis, as well as theories for its existence. Reproducibility shortcomings can arise at any stage, from funding to study publication.

Improving reproducibility starts with the funding of orthopaedic research. If those in charge of funding do not require statements about incorporating transparency and reproducibility in a project, such efforts may not be included in proposals. Among authors, a lack of reproducibility may arise from poor training of research staff, which can undermine even the most carefully crafted methodology. Further, poor planning and preparation, such as not prospectively registering a study or failing to identify appropriate reporting guidelines to follow, can decrease reproducibility. Recently published research has shown that prospective registration and reporting guideline use are frequently omitted in high impact orthopaedic literature, with many top orthopaedic journals not requiring these methodological safeguards^10^. Furthermore, even the most accurate and meticulous work will be difficult to reproduce if data are not kept in a clean generalizable format that is easily shared with others. With regard to peer reviewers, if they do not hold the manuscripts to a high standard of transparent reporting and data/materials sharing, manuscripts will pass peer review without including the factors needed to increase the odds of reproducibility. Lastly, and perhaps most importantly, journal editors and publishers are the final gatekeepers for ensuring published research is both transparent and reproducible. If they do not require studies to have a high degree of methodological transparency and ease of reproducibility, funding agencies, authors, and peer reviewers are unlikely to make the extra effort to promote reproducible and transparent practices.

### Recommendations for Improving the Reproducibility and Transparency of Orthopaedic Research

#### Funding Agencies

Nearly 85% of biomedical research is estimated to be wasted due to correctable causes, resulting in tens of millions of dollars in wasted funding^18^. Consequently, some funding agencies have implemented safeguards to prevent research waste. For example, the National Institutes for Health Research, UK, has required the prospective registration of systematic reviews that it funds, which will improve both transparency and reproducibility of the work^19^. Similarly, orthopaedic funding agencies can take measures to selectively fund research geared toward reproducing prior research needing verification. In addition, these agencies can give priority to projects that include explicit statements about how the methodology was constructed to be reproducible and how appropriate materials, methods, and data sets will subsequently be provided to promote reproduction of results. Furthermore, these agencies could provide support for the development of appropriate training and outreach courses for orthopaedic researchers at both local and national meetings/conventions. Agencies should also provide access to checklists and guidelines for conducting reproducible research within grant application information materials so that authors can incorporate these standards into their methodology from the inception of the proposal. By doing so, orthopaedic funding agencies would save themselves, and the field of orthopaedics in general, time and money by ensuring the research has the utmost methodological quality and transparency and is easily reproducible by others. Similar efforts have already been made by the NIH^1^.

#### Primary Investigators

Proper preparation by primary investigators such as training research staff to be knowledgeable about reproducible research practices, proper prospective trial registration, reporting guideline adherence, and clean and generalizable reporting of data will set studies up for success with regard to reproducibility from inception. Primary investigators could accomplish these goals by prospectively evaluating the standards and training provided by the NIH^1^ and incorporating these measures and training materials into their laboratories.

#### Journals and Peer Reviewers

Orthopaedic journals have the opportunity to strongly influence the transparency and reproducibility of orthopaedic research. If orthopaedic journals require manuscripts to meet standards that increase the odds of the work being reproducible, authors will have to conform to these standards. However, this outcome cannot happen overnight or with a simple change in policy. To have the greatest effect on orthopaedic reproducibility, journals must make changes gradually prior to enacting policies requiring reproducible practices. To start, high impact orthopaedic journals should invite leaders in orthopaedic surgery to write a collaborative editorial on the importance of conducting reproducible research and how the reproducibility crisis affects the field. This editorial should be open access across multiple journals to increase knowledge about the reproducibility crisis among as many orthopaedic surgeons as possible. Next, journals could provide links and education on reproducibility through their website and social media or in a newsletter. This same information should be provided to all potential peer reviewers to ensure they (and the general readership) understand the crisis and have the tools to know how to determine if a study provides the necessary elements to be reproduced. Journals should also provide this information in their instructions for authors and request that peer reviewers consider the authors’ compliance with guidelines in their recommendations to accept/reject the work. Next, journals could implement a section for “reproducibility and transparency” into their manuscript submission portal; completion of this section would be optional for a period of time. In this section, authors would explain the measures they took to ensure their work was transparent and easily reproducible and could upload any materials, methods, and data sets they deemed relevant. The next and final step would be for journals to require completion of the above section for manuscript submission. We recently reported that journals that require prospective trial registration and use guidelines had greater adherence to these methodological safeguards^10^. It can be reasonably concluded that journals that require these standards would increase adherence to them and therefore the transparency and reproducibility of orthopaedic research.

### Study Limitations

Our study has some limitations. First, we were limited to the content of each publication, and contact with the authors may have provided more thorough information on each study’s reproducibility. Additionally, an analysis of publications in a time frame different from ours might produce different results. Lastly, because our analysis was specific to orthopaedic journals, the results may not be generalizable to all areas of research.

### Conclusion

Our study evaluated the state of reproducibility in the field of orthopaedic surgery found the overall status to be poor, with over 90% of publications not providing statements on material or data availability. Furthermore, few studies in our sample were prospectively registered. Reproducible research is not solely the responsibility of those conducting the work—reproducibility and transparency in orthopaedic surgery can be improved by efforts from funding agencies, authors, peer reviewers, and journals alike.

