## Supplemental Table 1 for "Can Orthopaedics become the Gold Standard for Reproducibility? A Roadmap to Success"

**Supplemental Table 1:** Additional reproducibility characteristics

| **Characteristics** | | **Variables** |
| --- | --- | --- |
|  | | **No. (%)** |
| **Type of study (n=286)** | No empirical | 71 (24.83) |
|  | Meta-analysis | 8 (2.80) |
|  | Commentary with analysis | 1 (0.35) |
|  | Cost effect/Decisional analysis | 0 (0.00) |
|  | Clinical trial | 45 (15.73) |
|  | Case study | 15 (5.24) |
|  | Case series | 18 (6.29) |
|  | Cohort | 23 (8.04) |
|  | Chart review | 46 (16.08) |
|  | Case control | 2 (0.70) |
|  | Survey | 10 (3.50) |
|  | Laboratory | 27 (9.44) |
|  | Multiple | 4 (1.40) |
|  | Other † | 16 (5.59) |
| **Material available (n=9)** | Personal or institutional | 1 (11.11) |
|  | Supplementary information hosted by the journal | 3 (33.33) |
|  | Online third party | 0 (0.00) |
|  | Upon request | 5 (55.56) |
|  | Yes, material could be accessed and downloaded | 2 (22.22) |
|  | No, material could not be accessed and downloaded | 7 (77.78) |
| **Data available (n=13)** | Personal or institutional | 1 (7.69) |
|  | Supplementary information hosted by the journal | 4 (30.77) |
|  | Online third party | 2 (15.38) |
|  | Upon request | 5 (38.46) |
|  | Other (c) | 1 (7.69) |
|  | Yes, data could be accessed and downloaded | 5 (38.46) |
|  | No, data could not be accessed and downloaded | 8 (61.54) |
| **Documented data files contain all raw material (n=5)** | Yes, data files were clearly documented | 4 (80.00) |
|  | No, data files were not clearly documented | 1 (20.00) |
|  | Yes, data files contain all raw data | 2 (40.00) |
|  | No, data files do not contain all raw data | 2 (40.00) |
|  | Unclear if all raw data was available | 1 (20.00) |
| **Pre-registration (n=6)** | Yes, pre-registration was accessible | 6 (100.00) |
|  | No, pre-registration was not accessible | 0 (0.00) |
|  | Pre-registered on *clinicaltrials.gov* | 5 (83.33) |
|  | Other ¶ | 1 (16.67) |
|  | Hypothesis was pre-registered | 3 (50.00) |
|  | Methods were pre-registered | 2 (33.33) |
|  | Analysis plan was pre-registered | 3 (50.00) |
| **Protocol available (n=2)** | Hypotheses was included in the protocol | 2 (100.00) |
|  | Methods were included in the protocol | 2 (100.00) |
|  | Analysis plan was included in the protocol | 1 (50.00) |
| **Analysis script available (n=0)** | Personal or institutional | 0 (0.00) |
|  | Supplementary information hosted by the journal | 0 (0.00) |
|  | Online third party | 0 (0.00) |
|  | Upon request | 0 (0.00) |
| † Other included cross-sectional (n=15) and computational study design (n=1).  ‡ one article provided a statement saying, "All data generated or analyzed during this study are included in this published article."  ¶ www.controlled-trials.com. | | |
